# Machine learning models in electronic health records can outperform conventional survival models for predicting patient mortality in coronary artery disease

**DOI:** 10.1101/256008

**Authors:** Andrew J. Steele, S. Aylin Cakiroglu, Anoop D. Shah, Spiros C. Denaxas, Harry Hemingway, Nicholas M. Luscombe

## Abstract

Prognostic modelling is important in clinical practice and epidemiology for patient management and research. Electronic health records (EHR) provide large quantities of data for such models, but conventional epidemiological approaches require significant researcher time to implement. Expert selection of variables, fine-tuning of variable transformations and interactions, and imputing missing values in datasets are time-consuming and could bias subsequent analysis, particularly given that missingness in EHR is both high, and may carry meaning.

Using a cohort of over 80,000 patients from the CALIBER programme, we performed a systematic comparison of several machine-learning approaches in EHR. We used Cox models and random survival forests with and without imputation on 27 expert-selected variables to predict all-cause mortality. We also used Cox models, random forests and elastic net regression on an extended dataset with 586 variables to build prognostic models and identify novel prognostic factors without prior expert input.

We observed that data-driven models used on an extended dataset can outperform conventional models for prognosis, without data preprocessing or imputing missing values, and with no need to scale or transform continuous data. An elastic net Cox regression based with 586 unimputed variables with continuous values discretised achieved a C-index of 0.801 (bootstrapped 95% CI 0.799 to 0.802), compared to 0.793 (0.791 to 0.794) for a traditional Cox model comprising 27 expert-selected variables with imputation for missing values.

We also found that data-driven models allow identification of novel prognostic variables; that the absence of values for particular variables carries meaning, and can have significant implications for prognosis; and that variables often have a nonlinear association with mortality, which discretised Cox models and random forests can elucidate.

This demonstrates that machine-learning approaches applied to raw EHR data can be used to build reliable models for use in research and clinical practice, and identify novel predictive variables and their effects to inform future research.

## Introduction

Advances in precision medicine will require increasingly individualised prognostic assessments for patients in order to guide appropriate therapy. Conventional statistical methods for prognostic modelling require significant human involvement in selection of prognostic variables (based on an understanding of disease aetiology), variable transformation and imputation, and model optimisation on a per-condition basis.

Electronic health records (EHR) contain large amounts of information about patients’ medical history including symptoms, examination findings, test results, prescriptions and procedures. The increasing quantity of data in EHR means that they are becoming more valuable for research [1–5]. Although EHR are a rich data source, many of the data items are collected in a non-systematic manner according to clinical need, so missingness is often high [6,7]. The population of patients with missing data may be systematically different depending on the reason that the data are missing [8]: tests may be omitted if the clinician judges they are not necessary [9], the patient refuses [10], or the patient fails to attend. Multiple imputation to handle missing data can be computationally intensive, and needs to include sufficient information about the reason for missingness to avoid bias [11].

### Conventional versus data-driven approaches to prognostic modelling

Conventional statistical models, with a priori expert selection of predictor variables [1, 12], have a number of potential shortcomings. First, they may be time-consuming to fit and require expert knowledge of the aetiology of a given condition [13]. Second, such models are unable to utilise the richness of EHR data; a recent meta-analysis [1] found that EHR studies used a median of just 27 variables, despite thousands potentially being available [5]. Third, parametric models rely on assumptions which may not be borne out in practice. For example Cox proportional hazards models assume that a change in a predictor variable is associated with a multiplicative response in the baseline hazard that is constant over time. Nonlinearity and interactions need to be built into models explicitly based on prior clinical knowledge. Finally, missing data need to be handled using a separate process such as multiple imputation. Given that most imputation techniques assume that data are missing at random [11], this may mean that their results are unreliable in this context. Imputation can also be time-consuming both for researchers, and computationally given the large cohorts available in EHR data. Categorisation of continuous variables can accommodate nonlinear relationships and allow the inclusion of missing data as an additional category [14], but may substantially increase the number of parameters in the model.

Machine-learning approaches include automatic variable selection techniques and non-parametric regression methods which can handle large numbers of predictors and may not require as many assumptions about the relationship between particular variables and outcomes of interest [15]. These have the potential to reduce the amount of human intervention required in fitting prognostic models [16–18]. For example, random survival forests [19–21] can accommodate nonlinearities and interactions between variables, and are not restricted to a common baseline hazard for all patients, avoiding the assumptions inherent in Cox proportional hazards models.

Previous studies have used machine learning on EHR data for tasks such as patient classification and diagnosis [22–24] or predicting future hospitalisation [25], but we are unaware of any systematic comparison of machine learning methods for predicting all-cause mortality in a large, richly characterised EHR cohort of patients with stable coronary artery disease.

In this study we compared different analytic approaches for predicting mortality, using a Cox model with 27 expert-selected variables as the reference [13]. We aimed to:

1. Compare continuous and discrete Cox models, random forests and elastic net regression for predicting all-cause mortality;
2. Compare methods for handling missing data, and data-driven versus expert variable selection;
3. Investigate whether data-driven approaches are able to identify novel variables and variable effects.

## Methods

### Data sources

We used a cohort of over 80,000 patients from the CALIBER programme [26]. CALIBER links 4 sources of electronic health data in England: primary care health records (coded diagnoses, clinical measurements, and prescriptions) from 244 general practices contributing to the Clinical Practice Research Datalink (CPRD); coded hospital discharges (Hospital Episode Statistics, HES); the Myocardial Ischemia National Audit Project (MINAP); and death registrations (ONS). CALIBER includes about 4% of the population of England [27] and is representative in terms of age, sex, ethnicity, and mortality [26].

### Patient population

Patients were eligible to enter the cohort after being registered with the general practice for a year. If they had pre-existing coronary artery disease (myocardial infarction [MI], unstable angina or stable angina) they entered the cohort on their eligibility date, otherwise they entered on the date of the first stable angina diagnosis after eligibility, or six months after the first acute coronary syndrome diagnosis (MI or unstable angina) after eligibility. The six-month delay was chosen to differentiate long-term prognosis from the high-risk period that typically follows acute coronary syndromes. For patients whose index diagnosis was MI, we used information in CPRD and MINAP to attempt to classify the type of MI as ST-elevation myocardial infarction (STEMI) or non–ST-elevation myocardial infarction (NSTEMI).

Diagnoses were identified in CPRD, HES, or MINAP records according to definitions in the CALIBER data portal (www.caliberresearch.org/portal). Stable angina was defined by angina diagnoses in CPRD (Read codes) and HES (ICD-10 codes), repeat prescriptions for nitrates, coronary revascularisation (Read codes in CPRD or OPCS-4 codes in HES), or ischaemia test results (CPRD). Acute coronary syndromes were defined in MINAP or diagnoses in CPRD and HES.

### Follow-up and endpoints

The primary endpoint was death from any cause. Patients were followed up until they died (as identified in ONS or CPRD) or transferred out of practice, or until the last data collection date of the practice.

### Prognostic factors

Expert-selected predictors [28] were extracted from primary care (CPRD) and coded hospital discharges (HES). These included cardiovascular risk factors (e.g. hypertension, diabetes, smoking, lipid profile), laboratory values, pre-existing diagnoses and prescribed medication. Small-area index of multiple deprivation (IMD) score was derived from the patient’s postcode. Table 1 shows all the expert-selected predictors used.

**Table 1:**
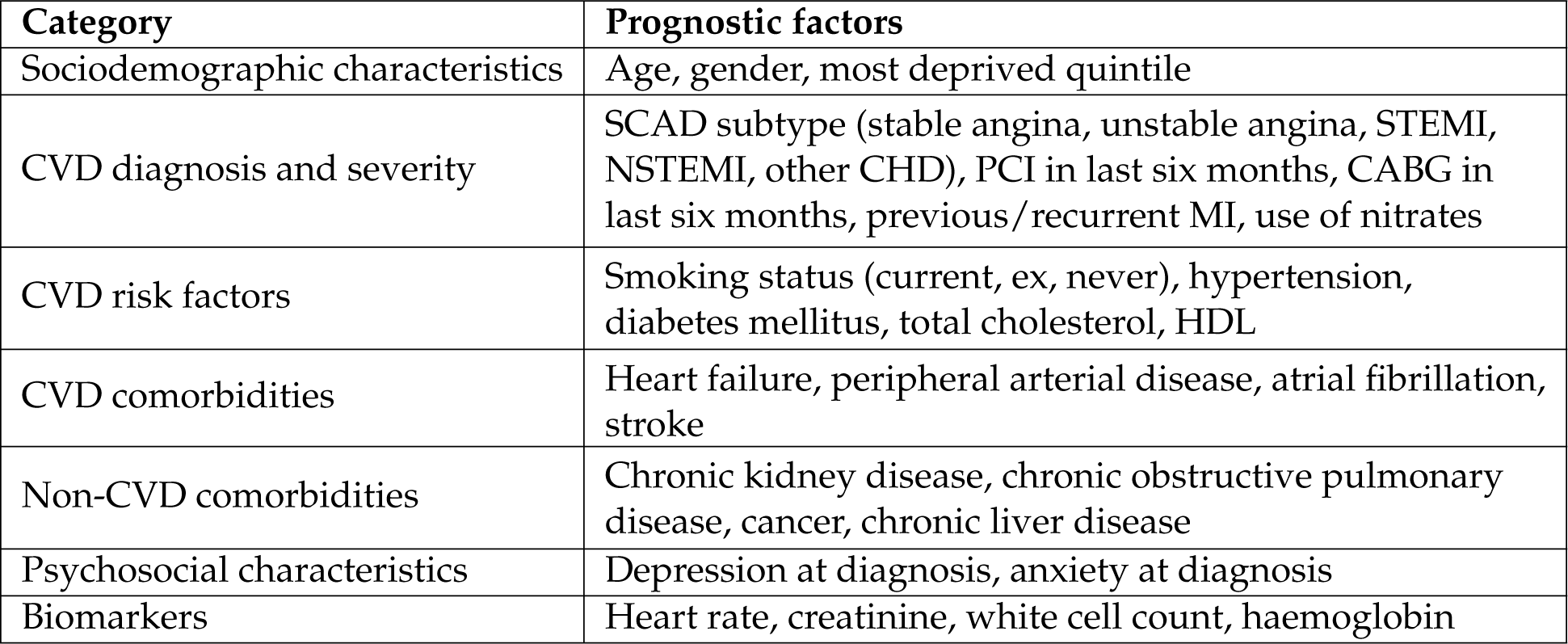
The 27 expert-selected predictors used. CVD = cardiovascular disease; PCI = percutaneous coronary intervention; CABG = coronary artery bypass graft; MI = myocardial infarction.

We also generated an extended set of predictor variables for the data-driven models, derived from rich data prior to the index date in CALIBER: primary care (CPRD) tables of coded diagnoses, clinical measurements and prescriptions; and hospital care (HES) tables of coded diagnoses and procedures. From each of these sources, we generated new binary variables for the presence or absence of a coded diagnosis or procedure, and numeric variables for laboratory or clinical measurements. We selected the 100 least missing variables from each source table, and the 100 least missing of both clinical history and measurements from the primary care data. After combining with the pre-selected predictors and removing duplicate and erroneous variables, there were 586 variables per patient in the extended dataset.

### Ethics

Approval was granted by the Independent Scientific Advisory Committee of the Medicines and Healthcare Products Regulatory Agency (protocol 14107) and the MINAP Academic Group.

### Statistical methods

#### Imputation of missing data

Multiple imputation was implemented using multivariate imputation by chained equations in the R package mice [29], using the full dataset of 115,305 patients. Imputation models included:

- Covariates at baseline, including all of the expert-selected predictors and additional variables: age, quadratic age, diabetes, smoking, systolic blood pressure, diastolic blood pressure, total cholesterol, HDL cholesterol, body mass index, serum creatinine, haemoglobin, total white blood cell count, CABG or PCI surgery in the six months prior to study entry, abdominal aortic aneurysm prior to study entry, index of multiple deprivation, ethnicity, hypertension diagnosis or medication prior to study entry, use of long acting nitrates prior to study entry, diabetes diagnosis prior to study entry, peripheral arterial disease prior to study entry, and history of myocardial infarction, depression, anxiety disorder, cancer, renal disease, liver disease, chronic obstructive pulmonary disease, atrial fibrillation, or stroke.
- Prior (between 1 and 2 years before study entry) and post (between 0 and 2 years after study entry) averages of continuous expert-selected covariates and other measurements: haemoglobin A1c (HbA1c), eGFR score [30], lymphocyte counts, neutrophil counts, eosinophil counts, monocyte counts, basophil counts, platelet counts, pulse pressure.
- The Nelson–Aalen hazard and the event status for all-cause mortality.

Since many of the continuous variables were non-normally distributed, all continuous variables were log-transformed for imputation and exponentiated back to their original scale for analysis. Imputed values were estimated separately for men and women, and five multiply imputed datasets were generated. We verified that the distributions of observed and imputed values of all variables were similarly distributed.

#### Training, validation and test sets

Before building prognostic models for all-cause mortality, we randomly split the cohort into a training set (
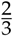
 of the patients), and a test set (the remaining 
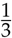
) which was used to evaluate performance of the final models. Performance statistics were calculated on repeated bootstrap samples from this held-out test set in order to estimate variability. Where cross-validation was performed other than elastic net regression, the training set was randomly split into three folds, and the model trained on two of the folds and performance validated against the third to establish optimal parameters, before those parameters were used to build a model on the full training set.

#### Continuous Cox proportional hazards models

Cox proportional hazards models are are survival models which assume all patients share a common baseline hazard function which is multiplied by a factor based on the values of various predictor variables for an individual.

Missing values in continuous variables were accommodated both by imputation, and, separately, by explicit inclusion in the model. The latter was done by first setting the value to 0, such that it did not contribute to the patient’s risk, and then a employing missingness indicator with an independent coefficient, meaning an additional binary dummy variable to account for whether a value is missing or not [8, 14]. For example, risk as a function of time *λ*(*t*) in a Cox model with a single continuous variable with some missingness would be expressed as

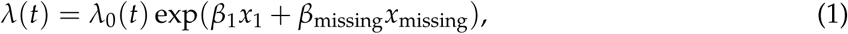

where *λ*_0_(*t*) is the baseline hazard function, *β*_1_ is the coefficient relating to the continuous variable; *x*_1_ is the variable’s value (standardised if necessary), set to 0 if the value is missing; *β*_missing_ is the coefficient for that value being missing; and *x*_missing_ is 0 if the value is present, and 1 if it is missing.

#### Missing categorical data

Missingness in categorical data can be dealt with very simply, by adding an extra category for ‘missing’. This is compatible with both Cox models (in which variables with *n* categories are parameterised as *n* − 1 dummy variables) and random forests (which can include categorical variables as-is).

#### Discrete Cox models for missing values

Another method to incorporate missing data is to discretise all continuous predictor variables, and include missing data as an additional category [8, 14]. Choice of category boundaries then becomes an additional hyperparameter.

Discretisation schemes were chosen by cross-validation to avoid overfitting. Discretising a value too finely may allow a model to fit the training data with spurious precision and generalise poorly to new data, whilst doing so too coarsely risks losing information carried by that variable. Every continuous variable *i* was split into 10 ≤ *n_i_* ≤ 20 bins, which were assigned in quantiles; for example, a 10-bin discretisation would split patients into deciles. An additional category was then assigned for missing values, resulting in *n_i_* + 1 coefficients per variable when fitting the Cox model.

#### Random survival forests

Random survival forests [19–21] is an alternative method for survival analysis which has previously been used to model deaths in the context of cardiovascular disease [31]. It is a machine-learning technique which builds a ‘forest’ of decision trees, each of which calculates patient outcomes by splitting them into groups with similar characteristics. These hundreds to thousands of decision trees each have random imperfections, meaning that whilst individual trees are then relatively poor predictors, the averaged result from the forest is more accurate and less prone to overfitting than an individual ‘perfect’ decision tree [32].

At each node in a decision tree, starting at its root, patients are split into two branches by looking at an individual covariate. The algorithm selects a split point which maximises the difference between the survival curves of patients in the two branches defined by that split. For a continuous parameter, this is a value where patients above are taken down one branch, and patients below are taken down the other; for a categorical variable, a subset of values is associated with each branch. Split points are defined so as to maximise the homogeneity within each branch, and the inhomogeneity between them.

Decision trees for regression or classification often use variance or Gini impurity, respectively. Survival forests, by contrast, maximise the difference between survival curves as measured by the logrank test [19], or optimise the C-index using which branch a patient falls into as a predictor [21]. After testing, we found splitting on C-index to be too computationally intensive for negligible performance benefit, so logrank tests were used.

This process is repeated until the leaves of the tree are reached. These are nodes where either there are no further criteria remains by which the patients at that leaf can be distinguished, including the possibility of only a single patient remaining, or splitting may be stopped early with a minimum node size or maximum tree depth.

In a random forest, each tree is created using only a random sample with replacement of the training data, and only a subset of parameters is considered when splitting at each node. A ‘forest’ comprising such trees can then use use the average, majority vote or combined survival curve, for regression, classification and survival forests, respectively, to aggregate the results of the individual trees and predict results for new data.

Random survival forests are thus, in principle, able to model arbitrarily shaped survival curves without assumptions such as proportional hazards, and can fit nonlinear responses to covariates, and arbitrary interactions between them.

One issue with random forests is that variables with a large number of different values offer many possible split points, which gives them more chances to provide the optimal split. One approach to combat this is to use multiple-comparisons corrections to account for the resulting increased likelihood of finding a split point in variables where there are many options [33, 34], but this is complicated by split statistics for adjacent split points being correlated. To avoid this issue, we instead used a value *n*_split_ to fix the number of split points tested in continuous variables which also has the advantage of reducing computational time for model building [35].

Many methods have been proposed for dealing with missing data in random forests, and these primarily divide into two categories. Some methods effectively perform on-the-fly imputation: for example the surrogate variable method, proposed in the classic classification and regression trees (CART) decision tree algorithm, attempts to use nonmissing values to find the variable which splits the data most similarly to the variable initially selected for the split [36]. Other methods allow missing values to continue through the network, for example by assigning them at random to a branch weighted by the ratio of nonmissing values between branches, known as adaptive tree imputation [19], or sending missing cases down both branches, but with their weights diminished by the ratio of nonmissing values between branches, as used in the C4.5 algorithm [37]. We used adaptive tree imputation, with multiple iterations of the imputation algorithm due to the large fraction of missing data in many variables used [19].

The number of iterations of the imputation process, number of trees, number of split points (*n*_split_), and number of variables tested at each split point (*m*_try_) were selected with cross-validation.

#### Variable selection

Performance can be improved by variable selection to isolate the most relevant predictors before fitting the final model. We evaluated various methods for variable selection: ranking by completeness (i.e. retaining those variables with the fewest missing values); by permutation variable importance (which measures a variable’s contribution to the final model performance by randomly shuffling values for that variable and observing the decrease in C-index), similar to varSelRF [38,39]; and by effect on survival, as measured by a logrank test between survival curves of different variable values (either true vs false, categories, or all quartiles for continuous variables). All of these are inherently univariate, and may neglect variables which have a large effect when in combination with others [15, 40].

After ranking, we fitted models to decreasing numbers of the highest-ranked variables and cross-validated to find the optimal number based on the C-index.

After selection, the contribution of variables to the final model was assessed by taking the permutation variable importance. There are a number of measures of variable importance in common use [41, 42], often using metrics which are specific to random forests, such as the number of times a variable is split upon, or the pureness of nodes following such a split. Permutation variable importance is more readily generalised: it is given by the reduction in performance when values of a particular variable are shuffled randomly within the dataset, thus removing its power to aid predictions—this allows it to be used to assess the importance of variables in other types of model, such as Cox models. The impact on model predictivity is assessed and, the larger the impact, the more important the variable is considered to be. Permutation variable importance was used in this study in order to allow comparisons across different types of model.

#### Elastic net regression

Elastic net regression extends regression techniques including Cox models to penalise complex models and thus implicitly select variables during the fitting procedure [40, 43]. It works by minimising the regularised error function

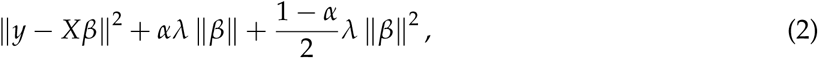

where *y* is the response variable, *X* are the input data, *β* is a vector of coefficients, 0 ≤ *α* ≤ 1 is a hyperparameter for adjusting the weight of the two different regularisation components, and *λ* is a hyperparameter adjusting the overall magnitude of regularisation. The hyperparameter *α* linearly interpolates between LASSO and ridge regression: a value *α* = 0 corresponds to pure LASSO, whilst *α* = 1 corresponds to pure ridge regression, with intermediate values representing a mixture of the two.

The value of *λ* is determined to some extent by the scale of the data, and so normalisation between parameters is important before fitting. In our case, values were either binary true/false when referring to the presence or absence of particular codes in a patient’s medical history, or represented discretised continuous values with an additional ‘missing’ category, binarised into a one-hot vector for fitting. Thus, since all values are either 0 or 1, no normalisation was undertaken before performing the elastic net procedure.

Hyperparameters *α* and *λ* were chosen by grid search with ten-fold cross-validation over the training set. Having identified the optimal *α*, this value was used to fit models on bootstrapped samples of the training set to extract model parameters along with an estimate of their variation due to sampling.

#### Model performance

Model performance was assessed using both the C-index, and a calibration score derived from accuracy of five-year mortality risk predictions.

The C-index measures a model’s discrimination, its ability to discriminate low-risk cases from high-risk ones, by evaluating the probability of the model correctly predicting which of two randomly selected patients will die first. It ranges from 0.5 (chance) to 1 (perfect prediction). The C-index is widely used, but its sensitivity for distinguishing between models of different performance can be low because it is a rank-based measure [44].

In particular, the C-index is unable to assess the calibration of a model, which reflects the accuracy of absolute risk predictions made for individual patients. A well-calibrated model is particularly important in the context of a clinical risk score, where a treatment may be assigned based on whether a patient exceeds a given risk threshold [45]. This was determined by plotting patient outcome (which is binary: dead or alive, with patients censored before the date of interest excluded) against mortality risk predicted by the model [44]. Then, a locally smoothed regression line is calculated, and the area, *a*, between the regression line and the ideal line *y = x* provides a measure of model calibration. Since a smaller area is better, we define the calibration score as (1 − *a*) such that a better performance results in a higher score. This means that, like the C-index, it ranges from 0.5 (very poor) to 1 (perfect calibration).

#### Variable effects

The partial dependency of Cox models on each variable is given by *β_i_ x_i_*, where *β_i_* is the coefficient and *x_i_* the standardised value of a variable *i*. The associated 95% confidence interval is given by *β_i_ x_i_ ±* Δ*β_i_ x_i_* where Δ*β_i_* is the uncertainty on that coefficient. Both the risk and the confidence interval are therefore zero at the baseline value where *x_i_* = 0. In discrete Cox models, coefficients for a given bin *j* translate directly into an associated *β_ij_* value, with an 95% confidence interval *β_ij_* ± Δ*β_ij_*. The lowest bin is defined as the baseline, and consequently has no associated uncertainty.

Due to their complex structure, interrogation of variable effects in random forests is more involved than for Cox models. They were assessed with partial dependence plots [46] for each variable. These are calculated by drawing many patients at random, and then predicting survival for each of them using the random forest many times, holding all variables constant for each patient except the variable of interest whose value is swept through its possible values for each prediction. This gives a curve of risks which we then normalised by its average, and then averaged over the set of patients to give a response to that variable. Relative changes of this risk can therefore be compared between models, but absolute values are not directly comparable.

## Results

### Patient population

A summary of the patient population used in this study is shown in Table 2. We initially identified 115,305 patients with coronary disease in CALIBER and, after excluding patients based on criteria relating the timing of the diagnosis and follow-up, 82,197 patients remained in the cohort. Imputation models included all 115,305 patients in order to increase precision, with the exclusion flag as an auxiliary variable in imputation models.

**Table 2:** Cohort summary, including all expert-selected predictors. Values are quoted to two significant figures and may not sum due to rounding. Continuous values are summarised as median (95% confidence interval). IMD = index of multiple deprivation; PCI = percutaneous coronary intervention; CABG = coronary artery bypass graft; MI = myocardial infarction.

| Characteristic | Summary |
| --- | --- |
| Patient population | 82,197 |
| Age | 69 (43–90) years |
| Gender | 35,192 (43%) women, 47,005 (57%) men |
| Social deprivation (IMD score) | 16 (4.0–52), 0.24% missing |
| Diagnosis | 57% stable angina<br>12% unstable angina<br>1.8% coronary heart disease<br>2.8% ST-elevated MI<br>3.1% non-ST-elevated MI<br>23% MI not otherwise specified |
| PCI in last 6 months | 4.6% |
| CABG in last 6 months | 2.0% |
| Previous/recurrent MI | 29% |
| Use of nitrates | 27% |
| Smoking status | 18% current<br>33% ex-smoker<br>40% non-smoker<br>8.2% missing |
| Hypertension | 88% |
| Diabetes mellitus | 15% |
| Total cholesterol | 4.8 (2.8–7.7) mmol l<sup>-1</sup>, 64% missing |
| High density lipoprotein cholesterol | 1.3 (0.70–2.3) mmol l<sup>-1</sup>, 78% missing |
| Heart failure | 10% |
| Peripheral arterial disease | 7.0% |
| Atrial fibrillation | 12% |
| Stroke | 5.4% |
| Chronic kidney disease | 6.7% |
| Chronic obstructive pulmonary disease | 35% |
| Cancer | 8.0% |
| Chronic liver disease | 0.41% |
| Depression at diagnosis | 19% |
| Anxiety at diagnosis | 12% |
| Heart rate | 72 (48–108) bpm, 92% missing |
| Creatinine | 94 (60–190) µmol l<sup>-1</sup>, 64% missing |
| White cell count | 7.1 (4.5–11) ×10<sup>-9</sup> l<sup>-1</sup>, 75% missing |
| Haemoglobin | 14 (9.9–17) g dl<sup>-1</sup>, 77% missing |
| Status at endpoint | 18,930 (23%) dead, 63,267 (77%) censored |

### Age-based baseline

A baseline model with age as the only predictor attained a C-index of 0.74. To calculate the risk and assess calibration, the Kaplan–Meier estimator for patients of age *x* years in (a bootstrap sample of) the training set is used, resulting in a well-calibrated model with a calibration score of 0.935 (95% confidence interval: 0.924 to 0.944).

### Calibration and discrimination of models

A summary of the performance results obtained for modelling time to all-cause mortality using various statistical methods and data is shown in Fig. 1.

**Figure 1:**
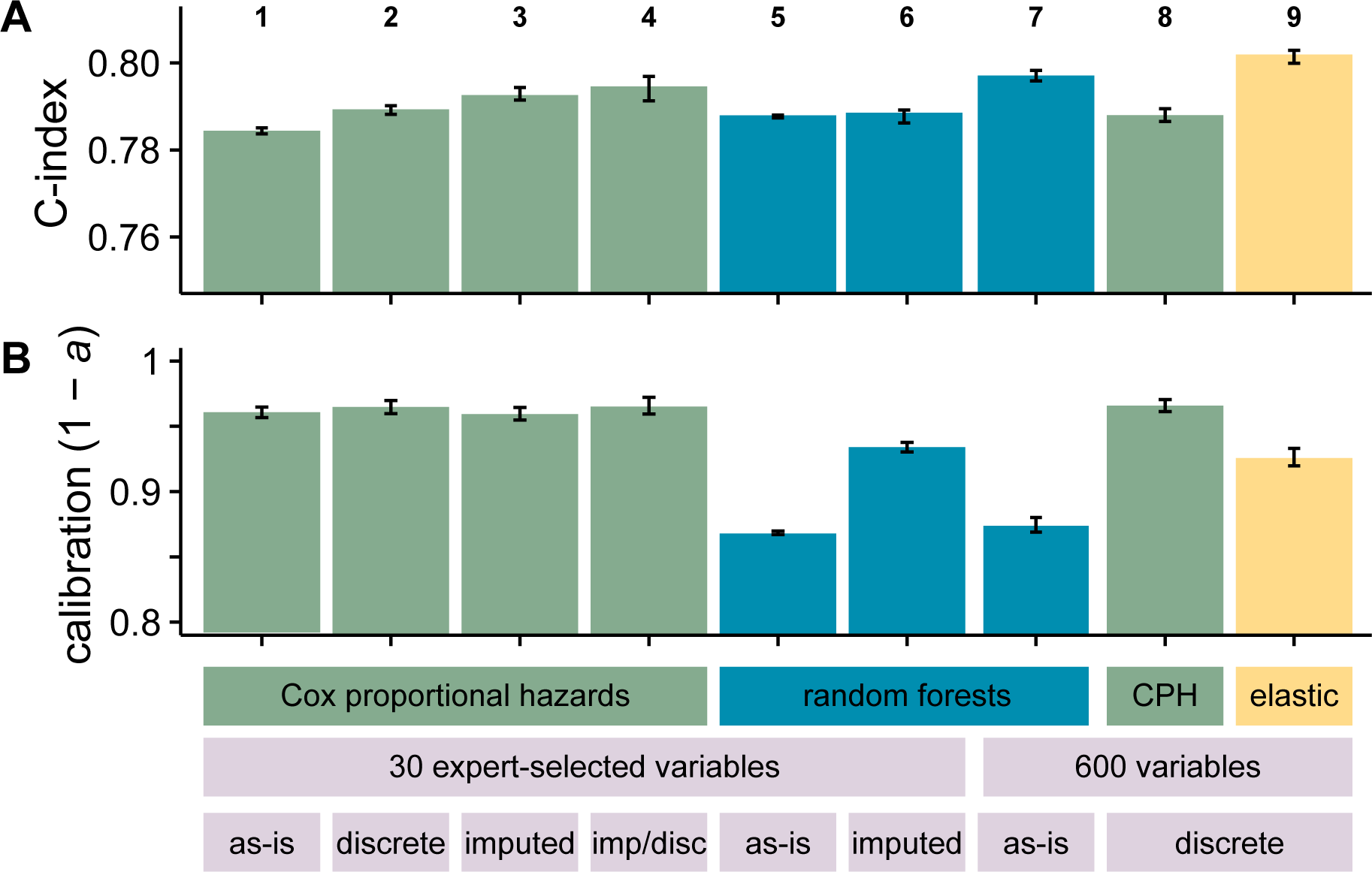
Performance measures for (A) discrimination (C-index) and (B) calibration (1 − *a*, where *a* is the area between the observed calibration curve and an idealised one for patient risk at five years) for the different modelling techniques used. Bar height indicates median performance across bootstrap replicates, with error bars representing 95% confidence intervals. Columns 1–4 represent variations on the Cox proportional hazards models using the 27 expert-selected variables used in Ref. [13]. Column 1 shows a model with missing values included with dummy variables; column 2 shows a model where continuous values have been discretised and missing values included as an additional category; column 3 shows a model where missing values have been imputed; and column 4 shows a model where missing values have been imputed and then all values discretised with the same scheme as column 2. Columns 5–7 show the performance of random survival forests. Column 5 uses the 27 expert-selected variables with missing values included with missingness indicators, and column 6 uses the imputed dataset. Columns 7 and 8 show models based on a subset of the 600 least missing variables across a number of different EHR data sources, selected by cross-validation. 7 is a random forest model with missing values left as-is, while 8 is a Cox proportional hazards model with continuous values discretised. Column 9 is an elastic net regression based on all 600 variables.

Comparing between models, the discrimination performance of random forests is similar to that of the best Cox models, but they performed worse in calibration, underperforming the Cox model with imputed data by 0.098 (0.105 to 0.091, 95% confidence interval on distribution of differences between bootstrap replicates). A calibration curve for this model is shown in Fig. 2(A). In view of this unexpectedly poor calibration performance, we also attempted to fit five-year survival with a classification forest. The binary outcome of being dead or alive at five years is essentially a classification problem, and it was possible that metrics for node purity during classification may be more reliable than those used in survival modelling. However, this proved to be both worse calibrated and worse discriminating than survival-based models, perhaps due to loss of the large number of censored patients in the dataset. A random survival forest implicitly takes censored patients into account by splitting based on logrank scores of survival curves, whereas a simple classification forest must discard them, perhaps losing valuable information.

**Figure 2:**
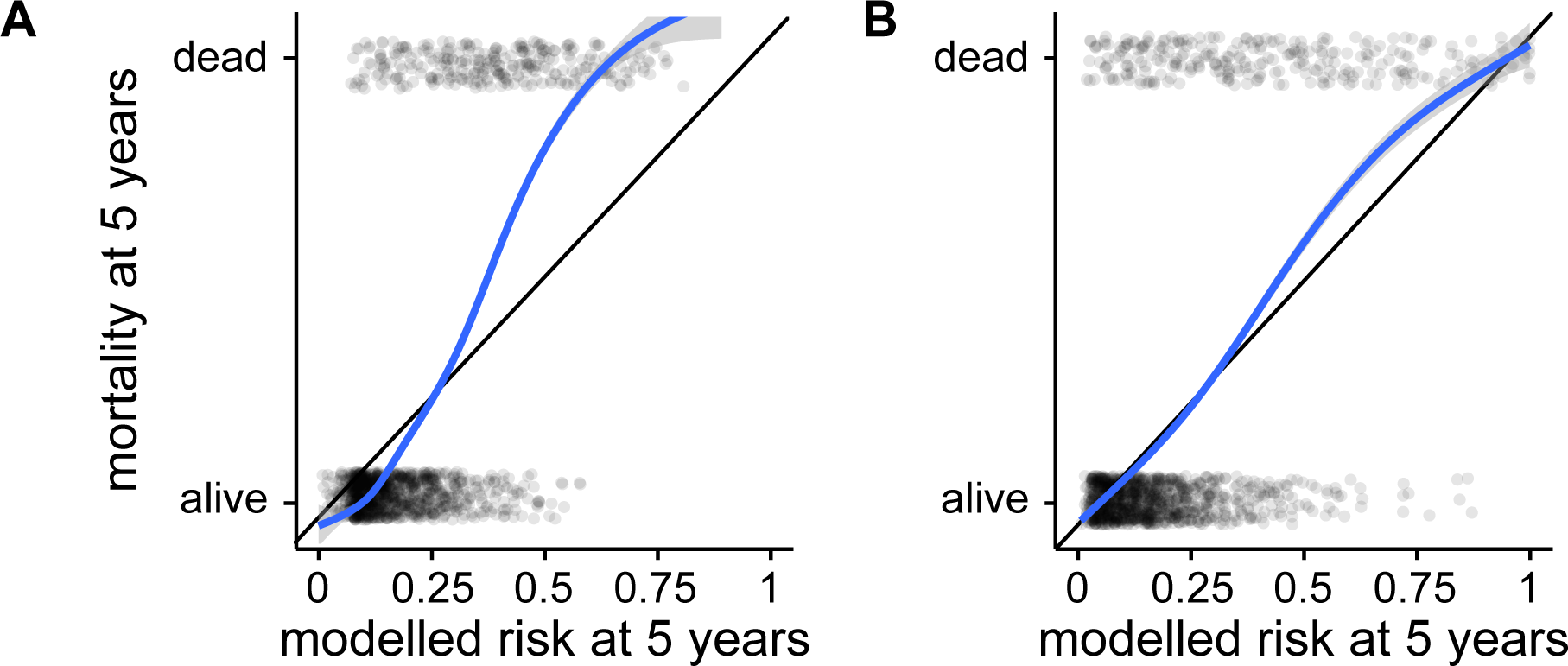
Example calibration curves for (A) model 5 (poorly-calibrated) and (B) model 9 (well-calibrated) from Fig. 1. Each semi-transparent dot represents a patient in a random sample of 2000 from the test set, the *x*-axis shows their risk of death at five years as predicted by the relevant model, and the *y*-axis shows whether each patient was in fact dead or alive at that time. The curves are smoothed calibration curves derived from these points, and the area between the calibration curve and the black line of perfect calibration is *a* in the calibration score, 1 − *a*.

Imputing missing values has some effect on model performance. Comparing the two continuous Cox models, column 1’s model includes missingness indicators and column 3’s model imputes missing values. Discrimination is very slightly improved in the imputed model, but calibration performance remains the same. Discretising continuous values without imputing has a similar effect, improving C-index slightly with little effect on calibration score.

To assess the impact of imputation on performance model-agnostically, we performed an empirical test by fitting a random forest model to the imputed dataset. The C-index remained similar, but calibration improves dramatically, almost achieving the same performance as the Cox model. This is suggestive but not decisive evidence that some information may leak between training and test sets, or from future observations, during imputation.

On the extended dataset, random forests fitted with all variables performed badly, even when cross-validated to optimise *m*_try_. After trying the different techniques discussed in Methods to rank important variables, by far the best random forest performance was achieved when variables were ranked by within-variable logrank test. This gave rise to a model using 98 of the variables, with a C-index of 0.797 (0.796 to 0.798). Unfortunately, even on the extended dataset random forests are similarly poorly calibrated to those fitted to the expert-selected dataset, under-performing the Cox model with imputed data by 0.086 (0.078 to 0.093).

A Cox model was also fitted to the extended dataset, with all continuous variables discretised into deciles. Variables were again ranked by within-variable logrank tests, and cross-validated to give a model comprising 155 variables. Its C-index performance was comparable to other models, and also has the highest median calibration performance of 0.966 (0.961 to 0.970).

Finally, an elastic net model was fitted to the discretised extended dataset. The optimal value of the hyperparameter *α* = 0.90 was determined by grid-search, and fixed for the remainder of the analysis. The hyperparameter *λ* was selected by ten-fold cross-validation on the full training dataset, returning nonzero coefficients for 270 variables. Bootstrapped replicates were again used to assess model performance, giving a calibration score of 0.075 (0.0679 to 0.0812), and the highest discrimination score of all models with a C-index of 0.801 (0.799 to 0.802). A calibration plot for this model is shown in Fig. 2(B).

### Effect of different methods for missing data

Fig. 3(A) shows coefficients associated with variables compared between a model based on imputation, and models based on accounting for missing values with missingness indicators, and discretisation. Most of the risks have similar values between models, showing that different methods of dealing with missing data preserve the relationships between variables and outcome.

**Figure 3:**
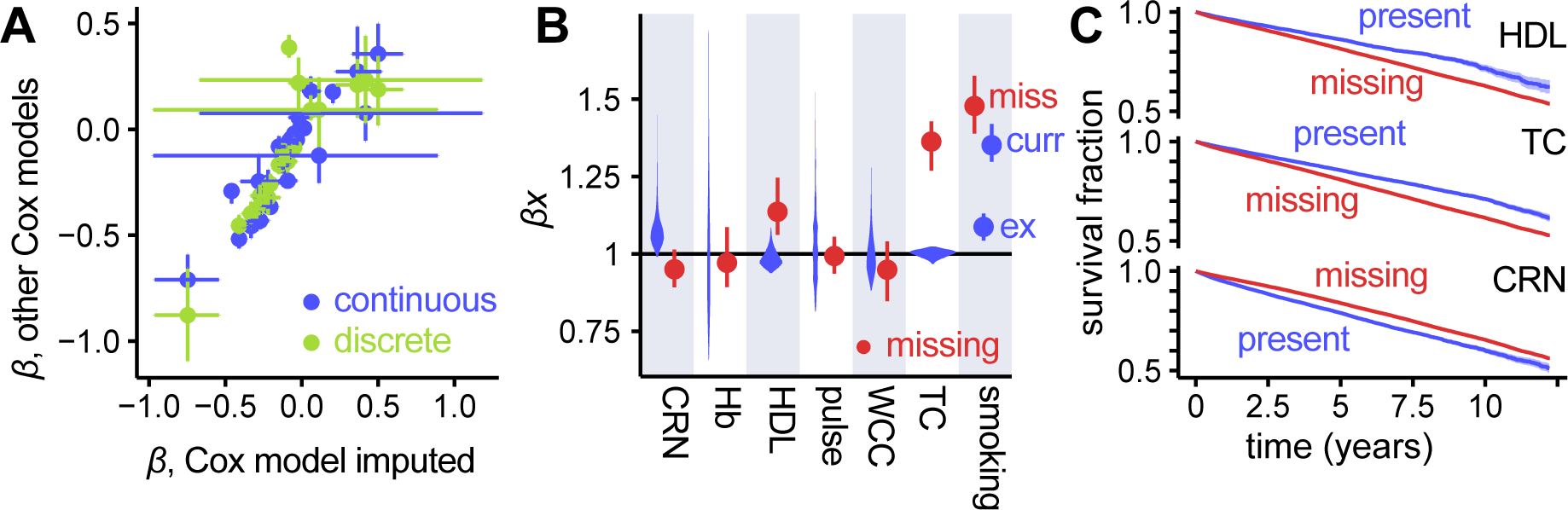
(A) Fitted coefficients for different Cox models compared. Values of risks from both the continuous model with missingness indicators, and the discretised model are plotted against the continuous imputed Cox model. There are fewer points for the discretised model as coefficients for continuous values are not directly comparable. The very large error bars on four of the points correspond to the risk for diagnoses of STEMI and NSTEMI. This is due to the majority (73%) of heart attack diagnoses being ‘MI (not otherwise specified)’ from which the more specific diagnoses were imputed, introducing significant uncertainty. (B) For the continuous Cox model with missingness indicators, risk ranges for the ranges of values for variables present in the dataset (violin plots) with risk associated with that value being missing (points with error bars). CRN = creatinine, HGB = haemoglobin, HDL = high-density lipoprotein, WBC = white blood cell count, TC = total cholesterol; for smoking status, miss = missing, ex = ex-smoker and curr = current smoker, with non-smokers as the baseline. (C) Survival curves for selected variables, comparing patients with a value recorded for that variable versus patients with a missing value. These can be compared with risks associated with a missing value, seen in (B): HDL and TC show increased risk where values are missing, whilst CRN shows the opposite, which is reflected in the survival curves.

Explicitly coding missing data as a missingness indicator allows us to associate a risk with that variable having no value in a patient’s record. These are examined in Fig. 3(B), compared to the range of risks implied by the range of values for a given variable. Having a missing value is associated with very different risk depending on which variable is being observed, indicating that missingness carries information about a patient’s prognosis. Where risks fall outside the range of risks associated with values found in the dataset, we validated these associations by plotting survival curves comparing patients with and without a particular value missing. These curves, shown in Fig. 3(C), agree with the coefficients shown in Fig. 3(B): variables whose missingness carries a higher risk than the normal range of values show a worse survival curve for patients with that value missing, and vice-versa.

### Variable and missing value effects

Next, we examined variable effects between models. These are shown in Fig. 4, with row (A) showing the continuous and discrete Cox models on unimputed data, and row (B) showing partial effects plots derived from the random survival forest models.

**Figure 4:**
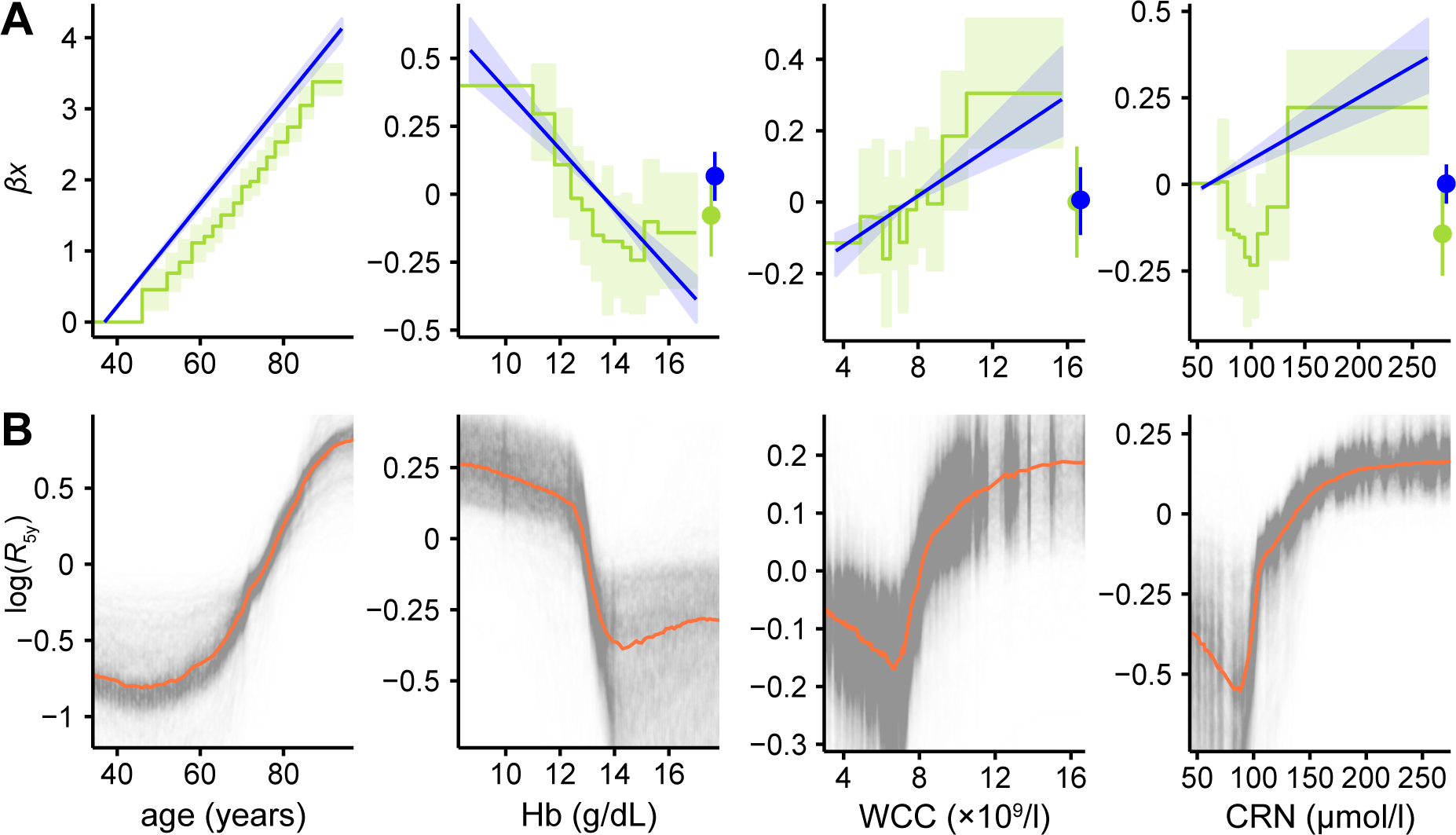
Variable effect plots from Cox models and partial dependence plots for random survival forests. (A) Comparisons between relative log-risks derived from the continuous Cox models (straight blue lines, light blue 95% CI region) against those derived from the discretised models (green steps, light green 95% CI region), together with log-risks associated with those values being missing (points with error bars on the right of plots). Confidence intervals on the continuous models represent Δ*β_i_ x_i_* for each variable *x_i_*, and hence increase from the value taken as the baseline where *x_i_* = 0. The lowest-valued bin of the discrete Cox model is taken to be the baseline and has no associated uncertainty. Discrete model lines are shifted on the *y*-axis to align their baseline values with the corresponding value in the continuous model to aid comparison; since these are relative risks, vertical alignment is arbitrary. (B) Partial dependence plots [46] inferred from random forests. Semitransparent black lines show the change in log-risk of death after five years from sweeping across possible values of the variable of interest whilst holding other variables constant. Results are normalised to average mortality across all values of this variable. Thick orange lines show the median of the 1000 replicates, indicating the average response in log-risk to changing this variable.

The partial effects of these variables show varying levels of agreement depending on model used. Risk of death rises exponentially with age, with the discrete Cox model’s response having a very similar gradient to the continuous model. Haemoglobin and total white blood cell count are in close agreement between these models too, with only slight deviations from linearity in the discrete case. Creatinine shows a pronounced increase in mortality at both low and high levels in the discrete model, which the linear model is unable to capture. The risks associated with having a missing value show broad agreement between the Cox models.

Random forest variable effect plots differ somewhat from the equivalent Cox model coefficients. The sign of the overall relationship is always the same, but all display quite pronounced deviations from linearity, especially at extreme values.

### Variable selection and importance in data-driven models

Fig. 5 shows the permutation variable importances for the 20 most important variables in the random forest and discrete Cox models fitted to the large dataset, after variable selection. All three models identify several variables which are also present in the expert-selected dataset, including the two most important (age and smoking status). There is also significant agreement within the 20 selected variables between models, with the top three appearing in the same order in the random forest and Cox model, and all appearing in the top five for the elastic net.

**Figure 5:**
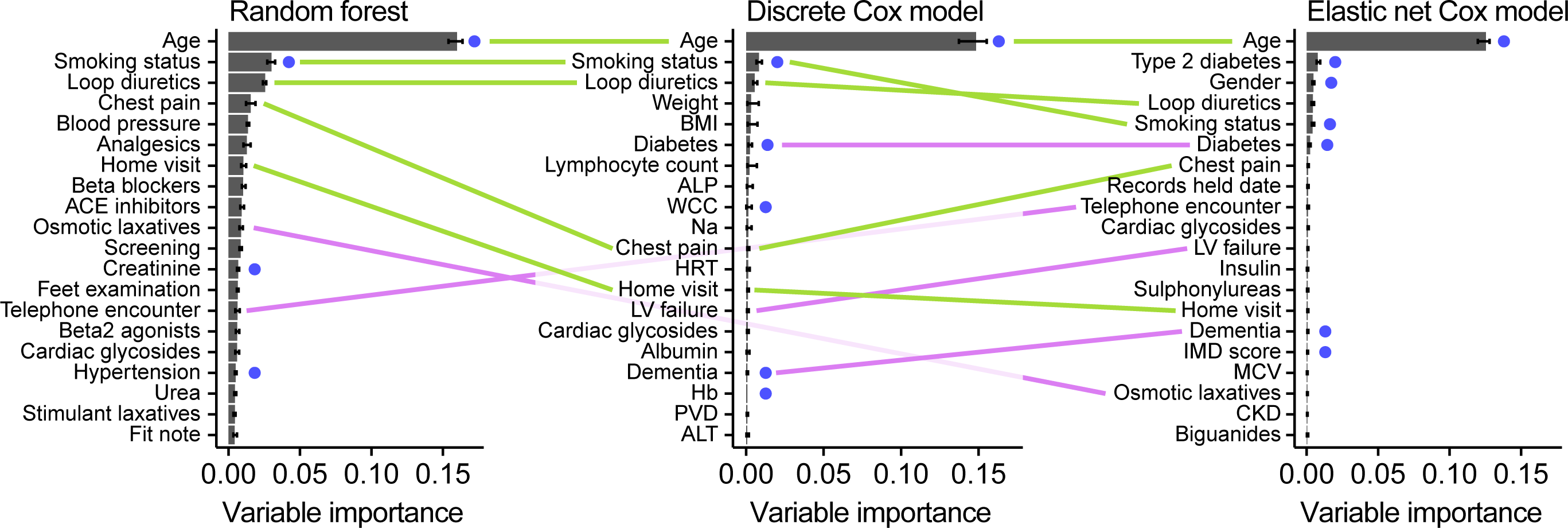
Top 20 variables by permutation importance for the three data-driven models, using random survival forests, discrete Cox modelling and elastic net regression. Variables which are either identical or very similar to those found in the expert-selected dataset are highlighted with a blue dot; variables appearing in two of the models are joined by a pink line (with reduced opacity for those which pass behind the middle graph); whilst variables appearing in all three are joined by green lines. Some variable names have been abbreviated for space: ACE inhibitors = angiotensin-converting enzyme inhibitors; ALP = alkaline phosphatase; analgesics = non-opioid and compound analgesics; ALT = alanine aminotransferase; Beta2 agonists = selective beta2 agonists; blood pressure = diastolic blood pressure; BMI = body mass index; CKD = chronic kidney disease; Insulin = intermediate‐ and long-acting insulins; LV failure = left ventricular failure; Hb = haemoglobin; MCV = mean corpuscular volume; Na = sodium; PVD = peripheral vascular disease; Records held date = date records held from; WCC = total white blood cell count.

When performing cross-validation to select variables for the Cox and random forest models, we observed large performance differences between ranking methods. Random forest variable importance on the full dataset performed poorly; it was a noisy measure of a variable’s usefulness, with significantly different variable rankings were returned on successive runs of the algorithm. Ranking variables by missingness performed better, but still significantly below the models using expert-selected variables. By far the best performance was achieved when variables were ranked by within-variable logrank test, which was used for both the random forest and discrete Cox models fitted here.

## Discussion

We compared Cox modelling and data-driven approaches for all-cause mortality prediction in a cohort of over 80,000 patients in an electronic health record dataset. All models tested had good performance. This suggests that, quantified by discrimination and calibration performance, there is no need to impute EHR data before fitting predictive models, nor to select variables manually. It also invites the use of data-driven models in other contexts where important predictor variables are not known.

### Advantages of data-driven approaches

Data-driven modelling has both practical and theoretical advantages over conventional survival modelling [5, 24]. It may require less user input, through automated variable selection. Further, both our method of converting continuous to categorical data (by quantiles), and random forest modelling are robust to arbitrary monotonic transformations of a variable, removing the need to manually normalise or transform data before analysis.

The ability to analyse data without the need for prior imputation also saves researcher time, and allows the effect of missingness in particular variables to be examined. It also has the potential to improve model performance in settings where missingness carries meaning. It is also possible for new patients to be assessed without the need for a clinician to impute, or take additional measurements, in order to obtain a risk score.

Discretisation of continuous values can accommodate nonlinear and nonmonotonic responses to variables, without requiring time-consuming choice and tuning of these relationships [47].

Machine learning techniques can also make use of more data, which in other datasets or settings may give rise to further improved performance over conventional models. Variable selection allows its use in contexts where risk factors are unknown.

### Comparison of modelling methods

We found that random forests did not outperform Cox models despite their inherent ability to accommodate nonlinearities and interactions [48, 49]. Random forests have a number of shortcomings which may explain this. First, only a random subset of variables (*m*_try_) are tried at each split, so datasets that contain a large proportion of uninformative ‘noise’ variables may cause informative variables to be overlooked by chance at many splits. Increasing *m*_try_ can improve performance, but often at a large cost in computation time. Second, when random forests are used for prediction, the predictions are a weighted average of a subset of the data, and are biased away from the extremes [50]. This may partly explain their poor calibration.

We did not find that discretisation of continuous variables improved model performance, probably because the majority of these variables had associations with prognosis that were close to linear, and the small improvement in fit was offset by the large increase in the number of model parameters.

Elastic nets achieved the highest discrimination performance of any model tested, demonstrating the ability of regularisation to select relevant variables and optimise model coefficients in an EHR context.

### Missing data

We also found that missing values may be associated with greater or lesser risk than any of the measured values depending on the potential reasons for a value’s presence or absence. For example, serum creatinine needs to be monitored in people on certain medication, and is less likely to be measured in healthy people. Conversely, missing high-density lipoprotein (HDL) or total cholesterol (TC), used for cardiovascular risk prediction in preventive care, was associated with worse outcomes.

There was little difference in model performance with or without imputation of missing values. It is possible that this did not have a large effect in the expert-selected dataset because only six variables carried missing values and, of these, only three had risks significantly different from the range of measured values. The majority of the variables were Boolean (e.g. presence of diagnoses) and were assumed to be completely recorded, where absence of a record was interpreted as absence of the condition.

One shortcoming of the use of imputation in our analysis is that, to our knowledge, no software is able to build an imputation model on a training set and then apply that model to new data. As a consequence, we performed imputation across the whole dataset, but this violates the principle of keeping training and test data entirely separate to prevent information ‘leaking’ between them. Further, because future values are used in the imputation process, this adds additional potential for introducing bias to models using such data. We investigated this empirically by fitting a random forest model to the imputed dataset (see Results) and found some evidence that bias may be introduced. If this is the case, the benefits of imputation to model performance may be less than suggested here.

### Variable selection

Finally, we found that data-driven modelling with large numbers of variables is sensitive to the modelling and variable selection techniques used. Random forests without variable selection performed poorly, which is evident from both the poor performance of fitted models, and the lack of utility of random forest variable importance as a measure by which to rank variables during selection.

Variable selection using the least missing data performed better, but not as well as expert selection. This may be because many of the variables with low missingness were uninformative. Variable selection using univariate logrank tests was far more successful, allowing models with slightly higher performance than expert selection, and discrete Cox models based on this displayed the best calibration performance.

Elastic net regression offered the best C-index performance, by fitting a Cox model with implicit variable selection. Since this selection process operates simultaneously across coefficients for all variables, it is possible that this explains its improved performance over ranking by univariate measures applied to random forests.

High-ranking variables in our final model (Fig. 5) seem plausible. Well-known prognostic factors such as age and smoking status were strongly associated with mortality, as expected. Many of the other variables identified, such as prescriptions for cardiovascular medications, are proxy indicators of the presence and severity of cardiovascular problems. Finally, some variables are proxies for generalised frailty, such as laxative prescriptions, home visits etc. These may be unlikely to considered for a prognostic model constructed by experts as they are not obviously related to cardiovascular mortality, and this demonstrates a potential benefit of data-driven modelling in EHR to both identify these novel variables, and improve model performance by incorporating them.

The important caveat is that these correlative relationships are not necessarily causal, and may not generalise beyond the study population or EHR system in which they were originally derived [1]. For example, our finding that home visits are a highly ranked variable is likely to be indicative of frailty, but its contribution will change depending on the criteria for such visits and the way they are recorded, which may vary between healthcare systems or over time.

### Disadvantages of data-driven approaches

Data-driven and machine learning based techniques come with several disadvantages. Firstly, the degree of automation in these methods is not yet complete; random forests worked showed poor performance on the full 586-variable dataset without variable selection. Use with other datasets or endpoints currently requires some researcher effort to identify optimal algorithms and variable selection methods. It would be useful to develop tools which would automate this, and test them across a multitude of different EHR systems and endpoints.

Secondly, it can be difficult to interpret the final models. Whilst it is possible to use variable effect plots to understand the relationship between a few variables and an outcome, this is prohibitively complex with high-dimensional models. In addition, not imputing missing values makes it difficult to interpret coefficients for partially observed variables, as they include a component of the reason for missingness. When prognostic models are used in clinical practice, it is important to be able to trust that the data are incorporated appropriately into the model so that clinicians can justify decisions based on the models.

Conventional statistical modelling techniques retain advantages and disadvantages which are the converse of these: models are more readily interpretable, and may generalise better, but at the expense of requiring significant expert input to construct, potentially not making use of the richness of available data, and only being applicable to complete data.

### Limitations of this study

These methods were compared in a single study dataset, and replication in other settings would be needed to demonstrate the generalisability of the findings.

The simple approach of using the top 100 least-missing variables from each table for development of the data-driven models is unlikely to be the optimal approach. In particular, some highly present variables are administrative and have little prognostic value, while significant but rare variables could be overlooked.

There are many types of model we did not consider in this study, including support vector machines and neural networks, which may provide improved prognostic performance. However, machine learning libraries for survival data are not well-developed, limiting the model types which can be tested without significant time devoted to software and model development.

### Conclusion

We have demonstrated that machine learning approaches on routine EHR data can achieve comparable or better performance than expert-selected, imputed data in manually optimised models for risk prediction. Our comparison of a range of machine learning algorithms found that elastic net regression performed best, with cross-validated variable selection based on logrank tests enabling Cox models and random forests to achieve comparable performance.

Data-driven methods have the potential to simplify model construction for researchers and allow novel epidemiologically relevant predictors to be identified, but achieving good performance with machine learning requires careful testing of the methods used. These approaches are also disease-agnostic, which invites their further use in conditions with less well-understood aetiology.

Eventually, machine learning approaches combined with EHR may make it feasible to produce fine-tuned, individualised prognostic models, which will be particularly valuable in patients with conditions or combinations of conditions which would be very difficult for conventional modelling approaches to capture.

## Supporting information

The scripts used for analysis in this paper are available at github.com/luslab/MLehealth. It is not possible to release the accompanying data due to patient confidentiality.

## Acknowledgements

This work was supported by the Francis Crick Institute which receives its core funding from Cancer Research UK (FC001110), the UK Medical Research Council (FC001110), and the Wellcome Trust (FC001110). NML and HH were supported by the Medical Research Council Medical Bioinformatics Award eMedLab (MR/L016311/1). The CALIBER programme was supported by the National Institute for Health Research (RP-PG-0407-10314, PI HH); Wellcome Trust (WT 086091/Z/08/Z, PI HH); the Medical Research Prognosis Research Strategy Partnership (G0902393/99558, PI HH) and the Farr Institute of Health Informatics Research, funded by the Medical Research Council (K006584/1, PI HH), in partnership with Arthritis Research UK, the British Heart Foundation, Cancer Research UK, the Economic and Social Research Council, the Engineering and Physical Sciences Research Council, the National Institute of Health Research, the National Institute for Social Care and Health Research (Welsh Assembly Government), the Chief Scientist Office (Scottish Government Health Directorates) and the Wellcome Trust.

